# Training data composition determines machine learning generalization and biological rule discovery

**DOI:** 10.1101/2024.06.17.599333

**Authors:** Eugen Ursu, Aygul Minnegalieva, Puneet Rawat, Maria Chernigovskaya, Robi Tacutu, Geir Kjetil Sandve, Philippe A. Robert, Victor Greiff

**Author notes:** E.U. and A.M. contributed equally to this work. To whom correspondence may be addressed. **Email:****, Preprint:** https://www.biorxiv.org/content/10.1101/2024.06.17.599333v2.abstract. **Author Contributions:** V.G. conceived the study. V.G., A.M., and E.U. designed analyses and visualizations. A.M., E.U. and P.A.R. performed analyses and visualizations. M.C. consulted on statistical matters. V.G., A.M., and E.U. wrote the first draft of the manuscript. All authors revised the manuscript and approved its contents. Given entirely equal contributions, A.M. and E.U. are free to list their respective names first in their respective CV. LLMs were used for text drafting. **Competing Interest Statement:** V.G. declares advisory board positions in aiNET GmbH, Enpicom B.V, Absci, Omniscope, and Diagonal Therapeutics. V.G. is a consultant for Adaptive Biosystems, Specifica Inc, Roche/Genentech, immunai, LabGenius, and FairJourney Biologics.

## Abstract

Supervised machine learning models depend on training datasets with positive and negative examples. Therefore, dataset composition directly impacts model performance and bias. Given the importance of machine learning for immunotherapeutic design, we examined how different negative class definitions affect model generalization and rule discovery for antibody-antigen binding. Using synthetic structure-based binding data, we evaluated models trained with various definitions of negative sets. Our findings reveal that high out-of-distribution performance can be achieved when the negative dataset contains more similar samples to the positive dataset despite a lower within-distribution performance. Furthermore, leveraging ground truth information, we show that binding rules discovered as associated with positive data change based on the negative data used. Validation on experimental data supported simulation-based observations. This work underscores the role of dataset composition, including negative data selection, in creating robust, generalizable, and biology-aware sequence-based ML models.

## Introduction

Supervised machine learning and deep learning models require the collection or construction of training datasets, which are composed of positive and negative examples (samples). Training datasets represent a fundamental part of a machine learning (ML) model’s success as they determine learned rules and prediction accuracy when applied to other tasks and domains (*out-of-distribution*, *OOD*) (Fig.1a) ^1–4^, and may introduce biases into the model’s decision-making process ^5,6^. Specifically, one important case of sample selection bias focuses on negative samples, that is, samples that do not represent the target class. It should be noted that in binary settings, negative and positive class definitions are mostly not independent. Negative sample bias has been discussed in different domains and contexts, such as text classification ^6–8^, protein-protein interaction prediction (e.g., immune receptor-antigen prediction) ^9–12^, and antimicrobial peptide prediction ^13^. While numerous strategies to address negative data sampling and data imbalance have been proposed ^14–16^, there is still a notable lack of research on how the negative dataset quality impacts the ML model’s outcome. More broadly, the influence of training set composition on generalization capacity and on the extraction of domain-specific knowledge (explainability) remains largely unexplored. In particular, within the sequence-level antibody-antigen binding problem, there is a wide array of qualitatively different negative datasets, offering an insightful terrain for evaluating the impact of negative data on model behavior.

**Fig.1.**
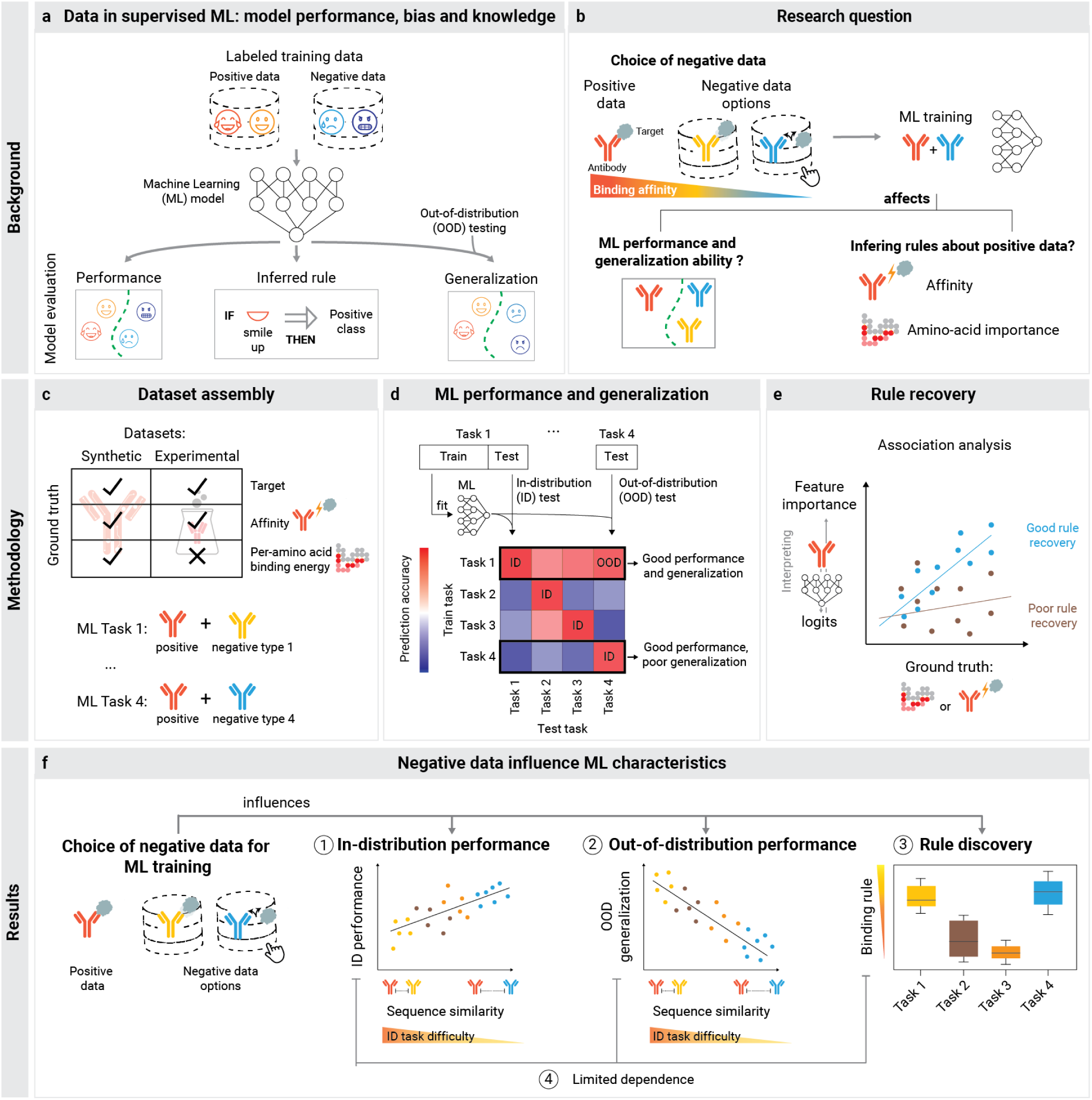
Training data composition determines machine learning generalization and biological rule discovery. **(a)** In supervised machine learning, particularly in binary classification, models are trained on labeled data divided into two categories: positive instances that align with the target concept and negative instances that do not. For illustration, positive emotion emojis represent the positive class, and all others represent the negative class. The standard way to evaluate an ML model is by testing it on a hold-out test dataset. Less frequently, the decision-making principles – rules – are examined, and even more rarely, the model’s generalization is assessed on an independent dataset where the data distribution may vary, referred to as an out-of-distribution (*OOD*) test. Notably, in cases where negative data is abundant and diverse, ML models can face significant challenges when tested with real-world data, which may exhibit shifts in the distribution of negative data. **(b)** Our research question explores how the selection of negative data for training the ML model affects its performance, generalization and rules in the context of antibody-antigen binding prediction. Specifically, we aimed to understand how negative data influences the identification of features associated with positive class antibodies. **(c)** Using both synthetic and experimental datasets with accessible ground truth information on affinity and target specificity, we constructed various binary classification tasks. These tasks shared the same positive data, while the negative data varied based on either affinity or target specificity. **(d)** After training the ML model on these tasks, we evaluated its performance on a hold-out test dataset, where the negative data had the same distribution as in training (in-distribution test). We assessed the model’s generalization ability by testing it on datasets with different distributions of negative data (out-of-distribution test). **(e)** The acquisition of affinity-related rules was estimated using explainability methods on trained models and comparing their outputs to ground truth information. **(f)** As a result, we found that the type of the negative dataset, and particularly its similarity to the positive dataset, correlates with both in-distribution performance and out-of-distribution generalization. Our results also revealed that negative data can either enhance or restrict the learning of affinity-based rules.

ML methods recently have demonstrated high performance in predicting immune receptor (antibody and T-cell receptor (TCR)) features such as their binding capability as well as pharmaceutically important biophysical properties ^17–24^ and, therefore, have become a promising approach in speeding up the development of immunotherapeutics. Regarding immunotherapeutics design, previous studies have demonstrated that the choice of negative data can affect prediction accuracy and generalization in antibody-antigen binding ^25–27^ and TCR specificity prediction ^12,28–31^. The influence of the negative dataset for antibody binding prediction was studied not only at the sequence level but also structurally, using simulated ^27^ and experimental data ^25^. Here, we explored the impact of training set composition on prediction accuracy and generalizability, with a particular emphasis on the discovery of binding rules (Fig.1b). We define learning of binding rules as identification of sequence-level binding energies and residue-level contributions to binding (energies).

The binding of immune receptors to their corresponding antigen is mainly determined by sequentially diverse complementarity-determining regions (CDRs) ^32,33^. Particularly, the CDR3 regions present in the heavy chain (CDRH3) of antibodies or beta chain (CDR3β) of TCRs with an average length of ∼16 amino acids contribute significantly to immune receptor-antigen binding ^32–34^. Therefore, in the current study, CDRH3 regions were used as a proxy to the full-length immune receptor sequences to train machine learning models. The three-dimensional interaction interface of the immune receptor-antigen complex is called a paratope on the immune receptor side and epitope on the antigen side. Existing state-of-the-art immune receptor-antigen binding databases contain <10^4^ data points ^35–40^ with known sequence/structures but, mostly, without the binding affinity of the paratope or epitope. To address this gap, synthetic data has emerged as a viable solution to benchmark machine learning methodologies ^1,41,42^. Synthetic data enables unconstrained access to ground truth data, enabling arbitrary composition of training datasets and, thus, a more controlled analysis of generalization and rule discovery. We have previously developed the antibody-antigen binding simulation framework “Absolut!” ^27^, enabling the generation of synthetic antibody-antigen complex structures at an unconstrained scale, where it was observed that accuracy-based rankings of ML methods trained on experimental data hold for ML methods trained on Absolut!-generated data, for both sequence- and structure-based tasks. These findings established the real-world relevance of ML benchmarking studies performed on Absolut!-generated data ^27^.

Here, we utilized the “Absolut!” framework ^27^ to create synthetic training datasets with various negative data compositions mimicking those commonly used and most readily available in immune receptor-antigen prediction studies (i.e., binders versus non-binders) (Fig.1c). We trained simple (to allow interpretability) neural networks on each type of dataset. We demonstrated that ML models generalized to unseen data of the same type. However, not all definitions of negative data in the training set led to models that could generalize to datasets containing different types of negative samples (Fig.1d,f). We also investigated the effect of dataset composition on the discovery of binding rules by using model explainability methods (namely DeepLIFT ^43–45)^ (Fig.1e). Explainability aims to reveal the reasoning underlying the model’s predictions, including which features and characteristics of the data have the most influence on those predictions. We show that biological rule discovery was a function of training dataset composition. We confirmed the conclusions of our work obtained on synthetic data using recently published “real-world”-relevant experimental large-scale antibody sequence data that was affinity-labeled for the HER2 breast cancer antigen ^46^ (Fig.1c).

Lastly, since our work relies on sequence-based data, its relevance reaches beyond the antibody and TCR field, to any sequence-based classification ^47^. This is especially useful in the case of sequence-based classifications where a sequence-to-fitness mapping that is relevant for learning can be defined. Our findings suggest that in such cases it is crucial to carefully consider the composition of ML training datasets to extract biological knowledge from sequence data.

## Results

## 1. Training dataset sequence composition influences prediction performance in ID and OOD binary classification tasks

### 1.1 Machine learning setup on synthetic and experimental data

We first investigated the influence of negative data composition on the performance and generalization of antibody-antigen binding predictors within the context of supervised binary classification. We define four distinct tasks and their corresponding datasets, with all datasets sharing the same positive CDRH3 sequences (binders), but with different types of negatives. These tasks broadly cover most scenarios of negative dataset definition (Table 1, Fig.2a). For a comprehensive evaluation of training dataset composition on ML performance, we employed synthetic CDRH3 sequences annotated with their binding energy to each of ten different antigens generated using the “Absolut!” framework ^27^. For each antigen, each of the four datasets shares the top 1% high-affinity percentile CDRH3 sequences as positives (Table 1). (i) The first dataset, “vs Non-binder”, contains the bottom 95% of low-affinity sequences to the same antigen, used as negative sequences; (ii) The second dataset, “vs Weak”, uses antibodies in the weak affinity range, from the top 1% to 5% percentile, as negatives. The “vs Non-binder” and “vs Weak” tasks are disjoint. (iii) The third dataset, “vs 1”, contains CDRH3 sequences that do not bind to the positive class antigen but that bind (top 1% affinity) to another predefined antigen. (iv) The “vs 9” dataset extends the “vs 1” by having binders to any other of the 9 antigens in equal amounts. Of note, datasets are balanced with 20 000 positive and 20 000 negative sequences, among which 10 000 are held-out as test data. We named the associated machine learning tasks by the name of the antigen (defining the positives) and the name of the dataset.

**Fig.2.**
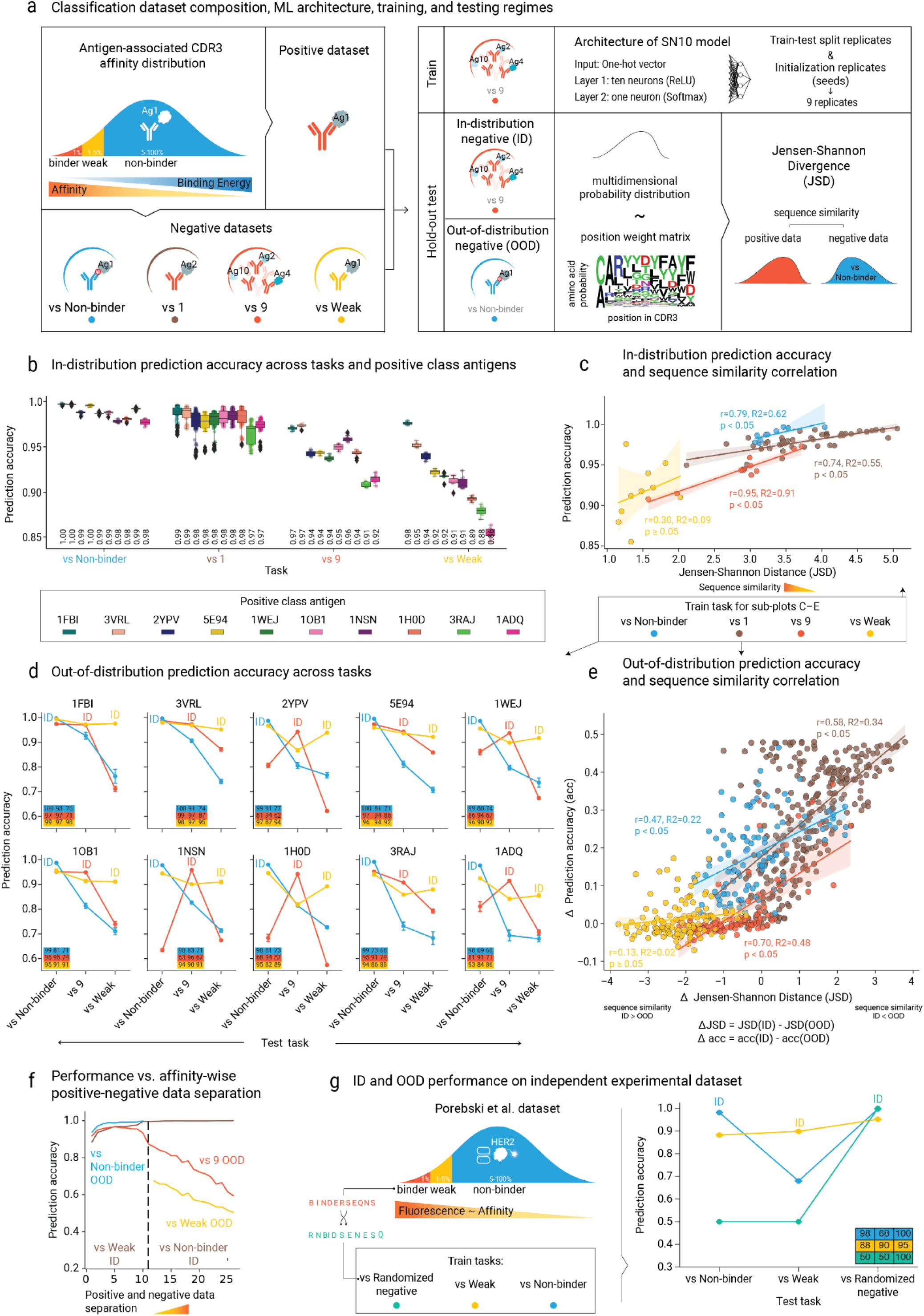
Classification performance varies across antigens, binding prediction tasks and the similarity between positive and negative data. **(a)** Antibodies related to a specific antigen can be categorized into affinity groups based on their binding energy: Binder, Weak, and Non-binder. The classification datasets are composed of a positive class of Binders to the target antigen and a negative class containing sequences that either bind to other antigen(s) (“vs 1” and “vs 9”), exhibit Weak binding (“vs Weak”), or are Non-binders (“vs Non-binder”) with respect to the same antigen (see Methods for details, Table 1). Low binding energy translates to high affinity and vice-versa. Each dataset was divided into train and test sets in five different ways (split replicates). We trained a machine learning model, SN10, consisting of a single layer with 10 neurons, employing four different initializations for one of the splits (seed replicates). To optimize computation time, we worked with nine replicates per task, which included one seed replicate combined with all split replicates and all seed replicates with one fixed split replicate. Evaluation was performed on the same type of task used during training – referred to as *in-distribution* (*ID*) testing, and on different tasks, termed *out-of-distribution* (*OOD*) testing. Additionally, we quantified the dissimilarity between the positive and negative classes in the test datasets using Jensen-Shannon distance (JSD) between the position-weight matrices of the corresponding sequences. **(b)** The *in-distribution* (*ID*) prediction accuracy of the SN10 model is grouped by negative dataset type and colored by positive class antigen. The boxes represent data between the 25th and 75th percentile. Numbers displayed indicate median prediction accuracies between replicates. **(c)** Relationship between the *ID* performance and the dissimilarity of the positive class and negative class sets of CDRH3 sequences (JSD). There are 10 data points for “vs 9”, “vs Non-binder” and “vs Weak” tasks and 45 points for “vs 1” task, each representing the average among replicates for respective combinations of positive and negative classes. The coefficient of determination (R2) and the Pearson correlation coefficient (r) were used to assess the relationship between Jensen-Shannon distance and prediction accuracy. Two-sided t-test p-values were below 0.05 for all tasks except for the “vs Weak” task. **(d)** *OOD* test performance. Each subplot corresponds to a positive class antigen. The hue indicates the training task (indicated by the “ID” label as well), and the x-axis reflects the test task. The plot includes tables that display the precise mean prediction accuracies. Error bars represent one standard deviation from the mean, capturing 68% of the variation in *OOD* performances. **(e)** The relationship between the change in Jensen-Shannon distance (JSD) values and the corresponding change in prediction accuracy (acc) between *ID* and *OOD* tasks. The ΔJSD and Δacc are calculated by taking the corresponding values for the *ID* test dataset and subtracting *OOD* test dataset values. The analysis includes 110 data points for the “vs 9”, “vs Non-binder”, and “vs Weak” tasks, and 270 points for the “vs 1” task. Two-sided t-test p-values were below 0.05 for all but the “vs Weak” task. **(f)** Change in *ID* and *OOD* performance as a function of positive–negative data separation in affinity space. The x-axis denotes negative data partitions, a higher partition number indicates a greater distance between positive and negative data in terms of affinity (see Supplementary Fig.10 for a schematic explanation). Partitions 0–10 correspond to “weak” negatives, while partitions 11–25 correspond to “non-binders”, a dashed vertical line marks the separation between them. The y-axis shows model performance: the brown line indicates *ID* performance, while the blue, red, and yellow lines show *OOD* performance on tasks defined in subfigure (a). The blue line (“vs Non-binder”) is shown only for partitions corresponding to the weak negatives (to the left of the dashed vertical line), and the yellow line (“vs Weak”) only for partitions corresponding to the non-binder negatives (to the right of the dashed vertical line). This subplot depicts results for 3VRL antigen. Similar plots for other antigens are shown in Supplementary Fig.10. **(g)** Results on Porebski et al. experimental dataset ^46^. The experimental dataset consists of CDRH3 sequences annotated with binding affinity, conceptually analogous to the synthetic dataset. The left panel illustrates the dataset generation strategy: the “vs Weak” and “vs Non-binder” datasets are defined identically to the “Absolut!” dataset (subplot **a**), while the “vs Randomized negative” dataset was created by randomly reordering the amino acids in the ‘Binder’ sequences. The right panel shows the SN10 test prediction accuracy for these datasets, where color denotes the training task and the x-axis represents the test task (similarly to subplot **d**). Data points are marked as ‘ID’ to indicate *in-distribution* performance; all other points represent *OOD* performance. Error bars signify one standard deviation from the mean. The inset table numerically presents mean *ID* and *OOD* prediction accuracies.

**Table 1.** Binary dataset composition and generation for binding prediction in adaptive immune receptors.

| Dataset type | Positive dataset | Negative dataset | Real-world relevance |
| --- | --- | --- | --- |
| vs Non-binder | Binder sequences with high affinity to the target antigen (range: 0–1% percentiles). | Sequences with low to no affinity (5–100% percentile) to positive class antigen – non-binders. | A common type of experimental dataset where binders and non-binders are identified by testing against a specific antigen and separated by a margin of binding affinity, ensuring that non-binders exhibit no considerable levels of binding. This dataset can be generated from experiments using fluorescent sorting or display selection. Alternatively, it can be acquired from broad-spectrum sequencing experiments without specificity information.<br>In the TCR field, it is similar to “negatives from background data”<sup>12</sup>. |
| vs Weak |  | Sequences with moderate affinity (1–5% percentile) to positive class antigen – weak binders. | A dataset where positive and negative data are close in affinity. It can be acquired through the same selection experiments mentioned above, but by selecting adjacent bins along the binding affinity gradient. |
| vs 1 |  | Sequences with high (affinity) binders to another antigen (range: 0-1% percentiles to another antigen). | A dataset consisting only of high-affinity binders to two antigens. Easily assemblable from public databases.<br>In the TCR field, it is similar to “negatives by shuffling” where the same epitopes and TCR from positive binding pairs are reused, but each TCR is paired with a different epitope<sup>12</sup>. |
| vs 9 |  | Sequences from high (affinity) binders of 9 | Assembled from individual binders to 10 antigens. It requires less data from each antigen and presumably |
|  |  | other antigens. | covers a wider landscape of negative data.<br>Also analogous to “negatives by shuffling” in the context of TCRs<sup>12</sup>. |

On each dataset, we trained a fully connected neural network with 10 hidden neurons and one neuron in the final layer (the model is hereafter termed SN10, see *Methods*). We utilized this relatively simple network architecture in our ML tasks as we previously showed that it outperforms classical ML models such as SVM and logistic regression, and more complex architectures such as convolutional neural networks or transformers do not yield any significant improvement in performance in similar ML tasks within “Absolut!” ^27^. To ensure the robustness of all results, the training process was replicated by varying both the train-test split composition and seed (influencing the model’s weights initialization, see Methods) (Fig.2a).

We repeated the analysis on synthetic data using recently published experimental data by Porebski et al. ^46^ that included affinity-labeled binding data for 24,790 CDRH3 sequences against HER2, further referred to as “the experimental dataset” (see Methods). In short, the authors developed an experimental method, which enables annotation of antibody sequence data with affinity-like information, replicating the data structure of the synthetic data we used.

### 1.2 ID prediction accuracy is a function of training dataset composition

To evaluate the capacity of trained models to learn generalizable rules, we compared their performance on *ID* test datasets, with the same positive/negative distribution as used in training (Fig.2a). Across all four types of *ID* test datasets, trained models achieved median prediction accuracies of over 0.85. Notably, we observed that the prediction accuracy decreased from the “vs Non-binder” task (range: 0.97–1.00, median: 0.99) to the “vs 1” task (range: 0.94–1.00, median: 0.98), followed by the “vs 9” (range: 0.91–0.98, median: 0.94), and finally the “vs Weak” task (range: 0.85–0.98, median: 0.92) (Fig.2b). Thus, the “vs Weak” task was the most challenging to achieve high accuracy on. Performance on shuffled control datasets, where sequence labels were randomly rearranged, reached as expected 0.5 (Supplementary Table 1). Thus, there are learnable CDRH3-based patterns that allow reaching high accuracies, which are lost due to label shuffling. In addition, we asked whether the SN10 model offers any advantage over logistic regression models (LR) and found that SN10 is on average 12% more accurate in more difficult tasks, such as “vs Weak” (Supplementary Fig.3c). This suggests that SN10 might exploit inter-positional dependencies, which are not covered by LR. Overall, SN10 and LR show a similar trend of performance across task types, with decreasing performance from “vs Non-binder” to “vs 1”, “vs 9” and “vs Weak”.

### 1.3 The distance between positive and negative datasets is linked to ID prediction accuracy

Given the gradation in *ID* performance across tasks, we asked whether the observed decrease in *ID* prediction accuracy may be attributed to the sequence dissimilarity between the positive and negative training datasets. We calculated the Jensen-Shannon distance (JSD) between the position-weight matrices (PWM) of the positive and negative class sequences (Fig.2a). The JSD exhibited an increasing trend from “vs Weak” to “vs 9”, then “vs 1” and “vs Non-binder”, suggesting increasing sequence divergence in this particular ordering. A corresponding increase in prediction accuracy accompanied this. Notably, the JSD between the positive and negative class sets demonstrated a stronger and significant (p<0.05) association with prediction accuracy for the “vs 9” (r=0.94), “vs 1” (r=0.73), and “vs Non-binder” (r=0.77) tasks, however not for the “vs Weak” task (r=0.30, p-value ≥ 0.05) (Fig.2c) and not for the shuffled controls (p-value ≥ 0.05 for all negative controls across each task type, Supplementary Table 1). The trend of increased prediction accuracy with higher JSD values remained largely consistent across all antigens (Supplementary Fig.3a). We excluded that the diversity of the overall training dataset, quantified by Shannon entropy, was significantly associated with *ID* prediction accuracy (p-value ≥ 0.05) (Supplementary Fig.1c). These results show that the SN10 model is more effective at discerning positive and negative training data when their sequence PWM-based Jensen-Shannon distance is greater. Furthermore, this suggests that binding prediction tasks with higher dissimilarity between positives and negatives are easier to learn when evaluated in an *ID* context.

### 1.4 OOD prediction accuracy is a function of training dataset composition

Having established that *ID* predictive performance depends on the task, we set out to investigate the *OOD* performance of trained models by taking models trained on one task and testing them on task types different then those of the training ones. For example, a model trained on the “vs 9” task would be tested using the “vs Non-binder” test data (Fig.2d, Supplementary Fig.3d). Expectedly, *OOD* performance of models was lower compared to the respective *ID* performance. The difference was highest for models trained on “vs Non-binder”, from 0.99 *ID* to 0.82 on the “vs 1” *OOD* task, 0.81 on the “vs 9” *OOD* and 0.72 on the “vs Weak” *OOD* tasks (Supplementary Fig.3d). The models trained on the “vs 9” task demonstrated the best *OOD* performance on the “vs 1” task, achieving an accuracy of 0.94. This result may be explained by the fact that “vs 9” includes binders to the same antigen as the “vs 1” task. Also, “vs 9” models exhibited generalization to the “vs Non-binder” separation task for 50% of antigens, resulting in a median accuracy of 0.91. We observed a high variation in the “vs 1” *OOD* performance depending on which other antigen was used to draw the negative sequences (Supplementary Fig.5). Generally, models trained on “vs 1” task demonstrated better *OOD* performance on the “vs 9” task, with an accuracy of 0.78, but performed relatively worse on the “vs Non-binder” task accuracy of 0.72 (Supplementary Fig.3d). The “vs Weak” task proved to be the most difficult *OOD* test task, as indicated by a median accuracy of 0.58 for models trained on “vs 1”, and 0.71 for models trained on “vs Non-binders” and “vs 9”. Importantly, models trained on the “vs Weak*”* task demonstrated the best overall generalization in the considered *OOD* tasks, with median prediction accuracies of 0.96 on the “vs Non-binder” task, 0.91 on the “vs 1” task, and 0.90 on the “vs 9” task although accuracies for certain antigens declined to 0.80 (Fig.2d). “Vs Weak” remains robust in both *ID* and *OOD* performance compared to other tasks, even when the number of negative data points in the test set increases (Supplementary Fig.11). When testing *OOD* generalization of logistic regression models, the same trend held, but overall, SN10 outperformed logistic regression in terms of *OOD* generalization (Supplementary Fig.3e). A negative control with shuffled labels ensured results were not due to chance: SN10 *OOD* on controls dropped to an accuracy of 0.5, (Supplementary Table 2).

We did not find a correlation between *ID* and *OOD* performance when considering all tasks together. However, when investigating *OOD* performances per task type, we observed associations between ID and OOD performance for “vs Weak” (Pearson r range: 0.8–0.98, p-value < 0.05) and “vs Non-binder” (Pearson r range: 0.37–0.57, p-value ≥ 0.05), however not for “vs 9” (Pearson r range: 0.09–0.11, p-value ≥ 0.05) (Supplementary Fig.4a).

In summary, dataset composition impacts the generalization capacity of models. Importantly, *ID* performance is not a reliable predictor of generalizability. This is exemplified by the “vs Non-binder” task. It yielded high *ID* prediction accuracies, but models trained on it yielded the lowest *OOD* accuracies. In contrast, the “vs Weak” task yielded the lowest *ID* accuracies but the best generalization ability.

### 1.5 OOD performance is associated with positive-negative similarity in training datasets

Next, we examined whether the stronger generalization ability of trained models to *OOD* tasks could be explained by greater similarity between the positive and negative datasets used during training. To address this, we analyzed similarity from two perspectives: sequence similarity and affinity-based distance.

First, we investigated the effect of sequence similarity on *OOD* performance by assessing the correlation between the difference in distances between negative and positive datasets in the training data (*ID*) and the corresponding *OOD* tasks. This difference was quantified as the gap in Jensen-Shannon distance between positive and negative data (ΔJSD = JSD(*ID* pos & neg) - JSD(*OOD* pos & neg)) and compared to the corresponding performance gap in prediction accuracy between *OOD* and *ID* tasks (Δacc = acc(*ID*) - acc(*OOD*)) (Fig.2e). We observed that models trained on “vs Weak” tasks had negative ΔJSD, which means that positive and negative data in the *OOD* test tasks were more distant compared to the *ID* task (“vs Weak”). “Vs Weak” models demonstrated relatively stable accuracy, and no correlation was observed between ΔJSD and Δacc (r = 0.13, p-value ≥ 0.05). Approximately half (52%) of Δacc values were less than or equal to zero. For the other training tasks (“vs Non-binder”, “vs 1”, “vs 9”), we observed a significant correlation between ΔJSD and Δacc ranging from 0.47 to 0.70 (p-value < 0.05). This indicates that if the positive and negative data in the training (and *ID* test) dataset are more distant than in the *OOD* test dataset, the *OOD* performance is likely to decrease. The extent of this decrease depends on the difference in the distance between positive and negative data in *ID* and *OOD* tests. However, the regression between ΔJSD and Δacc explained only 0.21–0.34 of the variance in the data, which may be partially due to the heterogeneity between antigens (Supplementary Fig.3b) but also suggests the existence of additional factors that contribute to model robustness. The shuffled controls corresponding to the *OOD* setting showed no significant correlation between ΔJSD and Δacc (Supplementary Table 2).

Second, we asked how the affinity-based similarity between positive and negative data affects *OOD* performance (Fig.2f, Supplementary Fig.10). Throughout this study, weak binders were defined as sequences in the 1st–5th percentile, and non-binders as those in the 5th–100th percentile of an antigen-specific binding energy distribution. Given that these are only two of many possible data partitions, we aimed to investigate to what extent prediction performance (*ID*, *OOD*) is a function of the distance in affinity space between binders (top 1% in affinity) and the negative data (lower than top 1%). To answer this question, we created 26 datasets with a gradually increasing affinity difference between positive and negative data. We trained SN10 models and evaluated the *OOD* performance on “vs 9” and, where relevant, on “vs Weak” and “vs Non-binder” tasks. Our findings show that *OOD* performance on “vs 9” deteriorates as the affinity gap between positive and negative data increases, with a similar trend observed in the “vs Weak” *OOD* task. Conversely, *OOD* performance on “vs Non-binder” improves, as increasing the affinity difference makes the training data more similar to the “vs Non-binder” task.

In summary, when models are trained on tasks with more distinct positive and negative datasets – as indicated by a larger JSD or affinity difference – they exhibit a decision boundary that, while effective within the *ID* context, may not be ideally configured for *OOD* tasks where the data distributions are closer. Conversely, training models on datasets characterized by lower JSD or affinity difference, where positive and negative data are inherently more similar, appears to foster a more adaptable and tighter decision boundary, leading to better generalization (*OOD* performance).

### 1.6 Experimental dataset replicates synthetic data findings on ID and OOD performance

We showed that negative data composition affects *ID* and *OOD* (Fig.2b,d) in synthetic datasets. To evaluate the persistence of this key finding from synthetic data, we leveraged a recently published experimental dataset ^46^, which resembles the “Absolut!” dataset as both contain CDRH3 sequences annotated with an affinity proxy measure (see Methods). We generated “vs Non-binder” and “vs Weak” tasks from the experimental dataset following the same strategy as in the synthetic data. Additionally, we created a “vs Randomized negative” task, where negative data was generated from positive binder sequences by randomly reordering the amino acids within each sequence while maintaining the global amino-acid composition. Thus, the ‘vs Randomized negative’ task requires the ML model to differentiate binder sequences from randomly shuffled ones, making it a relatively easy task. Finally, we analyzed the ID test-set performance and the *OOD* generalization performance of SN10 in these experimental datasets (Fig.2f).

Although the model trained on the “vs Randomized negative” dataset achieved 100% accuracy on its test set, in *OOD* scenarios, it failed to discriminate between classes (50% on “vs Non-binder”, 50% on “vs Weak”). Models trained on “vs Weak” and on “vs Non-binder” performed well on “vs Randomized negative”, with “vs Weak” achieving 95% accuracy compared to “vs Non-binder” (100%). Models trained on “vs Non-binder” had a higher recall, which contributed to their stronger performance in the ‘vs Randomized negative’ *OOD* task. Models trained on “vs Non-binder” also achieved high accuracy (98%) on their *ID* test, but could not perform well on “vs Weak” (68%). However, training on “vs Weak” resulted in comparable performance on both its *ID* test set (91%) and “vs Non-binder” *OOD* (88%). Therefore, the key finding on synthetic datasets, which showed that models trained on “vs Weak” tasks generalized best to *OOD* tasks, was recapitulated on the experimental data. Training on the “vs Weak” task likely results in models learning tighter decision boundaries, which might lead to more false-negative errors in both *ID* and *OOD* tests. In contrast, “vs Non-binder” has a looser decision boundary, allowing it to recall most of the positive data but also increasing the risk of false positives in more challenging *OOD* tasks.

To summarize, our results indicate that the composition of the negative dataset can affect predictive performance and generalization in binary classification tasks in synthetic and experimental data. Specifically, we found that predictive performance for *ID* and generalization capacity for *OOD* was associated with the similarity (affinity, sequence similarity) between positive and negative datasets. No clear association between *ID* and *OOD* was observed (Supplementary Fig.4a).

## 2. Training dataset composition determines the accuracy of biological rule recovery

Having shown that training negative data composition influences prediction accuracy and generalization on supervised binary sequence-based ML tasks, we set out to investigate whether negative data composition also impacts the capability to identify antibody-antigen binding rules. In this work, we define binding rules as either whole-sequence (per-sequence) binding energies or per-residue amino-acid-based contributions to binding energies. While additional binding rules may be defined, we focused on those for which comparable measures could potentially be extracted from ML models. Therefore, we acknowledge that our definition of binding rules is not exhaustive.

### 2.1 Training dataset composition impacts the learning of antibody binding energy

First, we investigated whether trained models have learned the energy properties of sequences beyond just classifying them as positives and negatives. Although there have been studies investigating the simultaneous prediction of binding status and affinity ^48,49^, binding status data is easier to generate experimentally. To investigate to what extent the SN10 model could recover a map from sequence to corresponding binding energy, we quantified the correlation of output logits and binding energy on a per-sequence level. Output logits are raw, pre-sigmoidal activation scores, measuring the model’s confidence in predicting the positive class (Fig.3a). An example of the 2D distribution of logit values and total binding energy is provided for the 3VRL antigen “vs Weak” task (Fig.3b inset).

**Fig.3.**
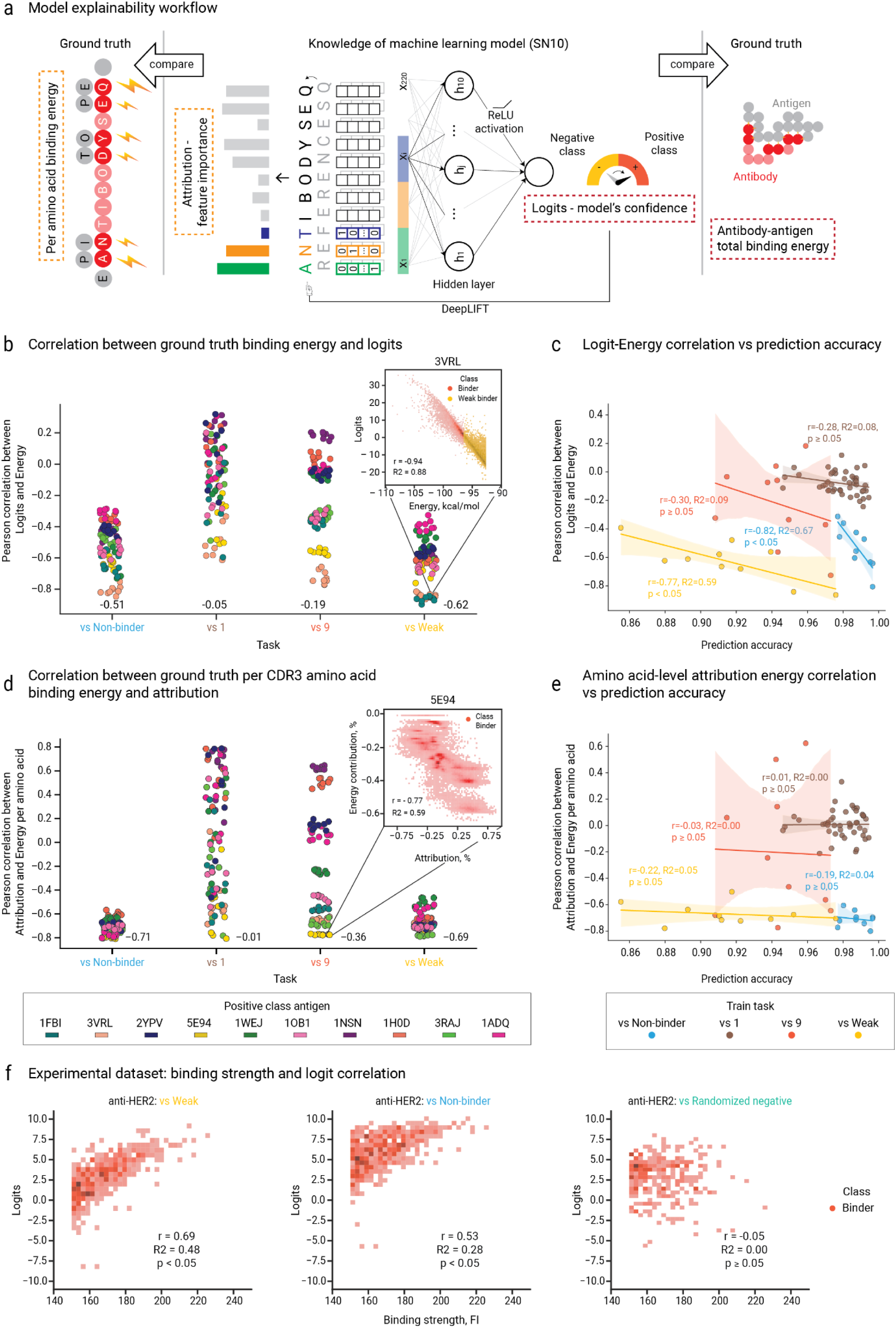
The choice of negative datasets can enhance the learning of binding rules of the positive class sequences. **(a)** To understand which antibody characteristics beyond their binding class models can acquire, we compared ground truth information available from “Absolut!” with the model’s interpretative quantities. As ground truth information, we used total antibody-antigen binding energy and per amino-acid binding energy. The model’s interpretive components were of two types: logits – the model’s output prior to the final activation, representing the model’s confidence in the predicted class; and feature attribution – the influence of a feature on logits. We employed the DeepLIFT attribution method to calculate how each feature in the input contributes to the prediction when compared to the reference sequence. In our study, we utilized DeepLIFT with ten distinct shuffles of the original sequence, subsequently averaging the results, as explained in the Methods section **(b)** Pearson correlation coefficients between logits and total binding energy in the positive class examples across different dataset types and antigens. We limited our analysis to the positive class examples from a dataset, since in practical scenarios, such as those corresponding to the “vs 1” and “vs 9” tasks, data regarding the binding of negative class sequences to the antigen of the positive class will be unavailable. The numbers at the bottom of the scatter plots indicate the medians of Pearson correlation coefficients within the tasks. Each dot in the scatter plots represents a correlation coefficient calculated for the relationship between logits and total binding energy values corresponding to a model-antigen pair. An example is provided for the “3VRL” antigen in the inset 2D-histogram (bins=100). **(c)** The relationship between SN10 model accuracy and logit-energy correlation coefficient was described for the preceding subplot. Pearson correlation coefficient (r) and coefficient of determination (R²) provide numerical descriptions of this relationship. The two-sided p-value is less than 0.05 for “vs Weak” and “vs Non-binder” tasks, and greater or equal for “vs 9” and “vs 1” tasks. For the tasks “vs 9”, “vs Non-binder”, and “vs Weak”, there are 10 data points in each, and for the “vs 1” task, there are 45 points. Each data point is an average across model train-test split and seed replicates. **(d)** Pearson correlation coefficients between energy contribution per CDRH3 amino acid and the SN10 amino acid attribution in the positive class samples across all dataset types and antigens. Numbers shown are median values. An example of a strong association is shown in the inset 2D histogram (bins = 100) for the 5E94 “vs Weak” task. The x-axis and y-axis values are expressed as signed percentages: each value represents the absolute attribution/energy for a specific amino acid in a CDRH3, divided by the sum of the absolute values of the attribution/energy for all amino acids in the CDRH3, with the sign preserved. **(e)** The relationship between SN10 model accuracy and per amino acid energy – attribution correlation coefficient, is described in the previous subplot. Similar to subplot (c). All p-values > 0.05. **(f)** Rule discovery efficiency by training task type, with all tasks constructed from the experimental Porebski et al. (28) dataset, as described in Fig.2f. A 2D histogram (30×30 bins) shows co-distribution of binding strength, measured as fluorescence signal intensity (FI), where higher fluorescence indicates stronger binding, and SN10 logit values for anti-HER2 scFv datasets. The Pearson correlation coefficient, indicated on each plot, is calculated solely for the sequences identified as binders.

Generally, “vs 1” and “vs 9” tasks did not allow for per-sequence energy rule learning, as indicated by correlations of −0.05 and −0.19, respectively (Fig.3b). The exceptions were the 3VRL (peach) and 5E94 (yellow) antigens, for which the models learned the total binding energy rule most effectively, irrespective of the negative data type. In contrast, negative datasets originating from “vs Weak” and “vs Non-binder” allowed the models’ intrinsic binding energy inference to improve for all antigens. This improvement resulted in a median correlation of −0.62 between antigens (range: −0.33 to −0.90) for models trained with “vs Weak” and a median correlation of −0.51 (range: −0.29 to −0.85) for models trained with “vs Non-binder” datasets (Fig.3b). Label-shuffled controls did not exhibit significant correlations (Supplementary Table 3). To summarize, we demonstrate that rule-learning on a per-sequence level depends on the type of negative dataset used, with significant variation in the Pearson correlation between output logits and binding energy observed across tasks (one-way ANOVA, p-value=8.7e-58) and positive antigen types (one-way ANOVA, p-value=4.6e-25).

To quantify whether this rule learning affects ML performance, we correlated the dependence of logit and binding energy with *ID* and *OOD*. For *ID*, we found significant correlations only for “vs Non-binder” (Pearson r = −0.83) and “vs Weak” (Pearson r = −0.77) (Fig.3c). Prediction accuracy in *OOD* tasks was significantly associated with logit-based rule discovery in all cases (Supplementary Fig.4b). This suggests that when the model learns the total binding energy rule, it performs better at identifying antigen binders across various types of negatives.

In summary, trained models can effectively learn to rank sequences by binding energies using solely binary-labeled data. The efficiency of this rule recovery depends on the type of negative data and the characteristics of the target antigen. Finally, learning of total binding energy is linked to variations in *ID* and *OOD* performance, reflecting both the influence of negative data choice and antigen-specific factors.

### 2.2 Training dataset composition impacts learning of position-wise contribution to the binding energy

Having established that the model implicitly acquired knowledge about binding energies for specific tasks and antigens on the whole-sequence level, we further aimed to investigate the model’s ability to acquire knowledge regarding the contribution of individual CDRH3 amino acid residues to binding — a crucial aspect of the binding process. To explore learning on the residue level, we obtained the energy contribution per amino acid to the binding energy from the “Absolut!” simulation suite ^27^. Our hypothesis was that if the model successfully acquired binding rules as amino acid contributions to binding, we will observe negative associations between the trained model’s attention towards specific amino acids and their contributions to binding energies. This is because lower binding energy corresponds to stronger binding, therefore, we expect residues with lower binding energy to have a greater influence in classifying a sequence as a binder. For the SN10 model, we analyzed attribution values per residue to measure the model’s attention towards specific amino acids using an explainable deep learning technique called DeepLIFT ^45^. The correlations with per-residue energies are computed for positive class CDRH3 sequences within each dataset (Fig.3d). An example 2D distribution of amino-acids attributions and energy is provided for 5E94 antigen “vs 9” task (Fig.3d inset).

Consistent with the learning of **per-sequence** binding energies, which were presented in the previous section, the rules of **per-residue** energies are learned better when models train with “vs Weak” and “vs Non-binder” negatives (Fig.3d). Median correlations were stronger for “vs Non-binder” (−0.71) and “vs Weak” (−0.69), compared to “vs 9” (−0.36) and “vs 1” (−0.01) tasks. For several antigens (3VRL - peach, 5E94 - yellow, 3RAJ - light green) correlations were high regardless of the type of negative data (Fig.3d). These antigens’ feature (amino-acids) attributions correlate well across different tasks: “vs Weak,” “vs Non-binder,” and “vs 9” (Supplementary Fig.8a). Showing that in rare cases only positive data is sufficient to learn residue-wise binding rules. Some antigens in “vs 1” and “vs 9” were characterized by strong positive correlations, breaking the biologically sound link between low binding energy and high binding strength. Notably, in the “vs 1” task, switching the roles of positive and negative datasets restored high negative correlation (e.g., 2YPV vs. 5E94, r=0.78 → 5E94 vs. 2YPV, r=-0.8; Supp. Fig. 6C). This demonstrates a potential bias in the ML model, where it tends to learn binding rules favoring the sequences of the “easier” antigen, irrespective of their assignment as positive or negative. For example, SN10 exhibited bias toward 5E94 in both cases: 5E94(pos) vs. 2YPV(neg) and 2YPV(pos) vs. 5E94(neg). To summarize, the choice of negative data significantly influences the learning of binding rules about positive data, as evidenced by varying degrees of correlation between attributions and per-residue energies across task types (one-way ANOVA, p-value=3.5e-43, Fig.3d). Shuffled-label controls exhibited no significant correlations (Supplementary Table 3).

We found no significant association between the prediction accuracy and the correlation between per-residue energy and attributions, across all datasets (Fig.3e), providing another example of a negative dataset-dependent property of a trained network that is not reflected by the *ID* prediction performance. *OOD* prediction accuracy, on the other hand, was associated with attribution-based rule discovery for all “vs 1” *OOD* scenarios and for all “vs 9” *OOD* tasks, except when tested on each other. No significant correlation between *OOD* prediction accuracy and per-residue rule discovery was observed for models trained on “vs Weak” and “vs Non-binder” (Supplementary Fig.4b). These findings indicate that the per-residue energy rule alone is not sufficient to ensure high *OOD* performance. Still, knowledge of per-residue energy imposes a constraint on *OOD*, as models that assign high importance to high-binding-energy residues (positive correlation) fail in *OOD* tasks, as observed for some models trained on “vs 1” and “vs 9”.

As a summary, the inclusion of “vs Weak” and “vs Non-binder” negatives seems to facilitate the model’s ability to learn per-residue binding energies, although in some cases, positive data alone may provide sufficient information for this purpose. While the per-residue rule is not the most critical factor for *ID* and *OOD* performance, learning it incorrectly—with reversed correlations—can lead to a substantial decrease in *OOD* performance.

### 2.3 Investigating the extent of additivity in learned rules

We observed that the attributions for “vs Weak” and “vs Non-binder” correlated highly (Supplementary Fig.8.). Given that their *OOD* performances (a potential correlate of rule learning) differed, we sought to delineate the nature of the learned rules by investigating and highlighting the differences in the proportion of learned additive versus non-additive rules. To this end, we investigated the associations between energies and weights of logistic regression models–representing learners of only additive rules–trained in identical settings on synthetic data. Logistic regression is fundamentally SN10 without the middle hidden layer, lacking feature interaction components while retaining global non-linearity. The weights of the logistic model can be similarly interpreted as the attributions of SN10, and namely as contributions to the logits. We showed that the correlations per-sequence (logits vs total energy) and per-residue (weights vs per-residue energies) are similar in magnitude to the ones based on SN10 (Supplementary Fig.7a,b), e.g., median per-sequence correlation for “vs Weak” in SN10 (−0.62) closely matched that of logistic regression (r=-0.58).

We also found that the weights of logistic regression and attributions of SN10, i.e., the feature importances, correlated to varying degrees, depending on the task and antigen type. The highest correlation was observed for the “vs Non-binder” task (median across antigens: 0.84), while “vs 9” and “vs Weak” exhibited median correlations of 0.60. However, the correlation range for “vs Weak” was wider (Supplementary Fig.7c).

These findings suggest that, in the “vs Non-binder” task, per-residue importances derived from SN10 align more closely with logistic regression, indicating that feature attributions in this case are primarily driven by linear combinations of features. In contrast, the deviations observed in the “vs Weak” task imply that SN10 feature attributions incorporate feature interactions. Notably, the superior performance of SN10 in specific tasks supports the hypothesis that SN10 captures more complex rules, leading to better performance than logistic regression, particularly for the “vs Weak” task (Supplementary Fig.3c).

### 2.4 Training dataset composition drives per sequence binding energy recovery from experimental data

We investigated whether the rule discovery conclusions on synthetic datasets would be replicated on experimental datasets by applying the same methodology to the Porebski experimental dataset ^46^, as in the previous section. Note that, due to the nature of the experimental data, we had access only to a proxy of per sequence binding energies, without any available proxy for per amino acid contributions, in contrast to the synthetic data. Therefore, we examined the associations between the logits of the models trained on the experimental dataset and the fluorescence-based binding intensity measure (further called intensity), as previously defined ^46^ (see Methods). We observed high correlations between model logits and intensity (“vs Weak” r=0.84, p-value < 0.05; “vs Non-binder” r=0.82, p-value < 0.05), in line with the results from the synthetic dataset (Fig.3f). The models trained on datasets with randomized negative sequences did not learn the same associations (“vs Randomized negative” r=-0.05), despite the 100% test set accuracy, which might suggest that the model trained on “vs Randomized negative” could achieve high accuracy in distinguishing between the positive and negative class examples by leveraging simpler rules unrelated to binding, such as ones based on amino-acid frequency. Although learning binding energies, which are continuous quantities, from binary classification was a non-trivial result, it has been recently shown in a different context ^50^.

## Discussion

The definition of “negative” and “positive” data is crucial in supervised machine learning settings, but how the definition of negative and positive data changes ML model behavior (prediction accuracy, generalization, rule inference) has not been thoroughly investigated. Here, we showed that *ID* and *OOD* performance, and the rule discovery vary as a function of training dataset composition in the antibody-antigen binding prediction task. As mentioned throughout the text, there is considerable discussion in the TCR-epitope prediction field about the choice of negative data ^12,28–31^. Given that TCR-epitope prediction settings are very similar to this work’s ML setup (supervised classification, several classes of sequences in the datasets), this work provides guidance on immune receptor dataset construction, benefiting both the BCR and TCR fields. More generally, our work shows that ML dataset construction is far from trivial and needs to be planned carefully for robust *ID* and *OOD* performance and rule discovery.

### ID performance does not predict OOD predictive performance

Briefly, we found that for both synthetic and experimental data, in the *ID* setting, prediction performance was high (with “vs Weak” having the lowest performance, Fig.2b) but varied across training data compositions. In contrast, in the *OOD* setting, only models trained on “vs Weak” performed consistently well (Fig.2d, Supplementary Fig.3, Supplementary Fig.11). Thus, in line with previous observations in an entirely different domain ^51,52^, we found a trade-off between *ID* and *OOD* settings across tasks. While within “vs 9” task *ID* performance did not predict *OOD* performance, “vs Non-binder” indicated a non-significant positive trend in the *ID-OOD* relationship, and “vs Weak” exhibited a positive correlation (Supplementary Fig.4a), suggesting that these ML models learned invariant (robust) features of the positive data, making *ID* informative of *OOD*. In cases of negative or no correlation, the model likely relied mostly on spurious features ^51^.

We found that sequence similarity (measured by JSD) between positive and negative data correlated with *ID* and *OOD* performance (Fig.2c,e). The smaller the distance (JSD), the lower was *ID* performace but the more stable *OOD* was. Similarly, when positive and negative data in the training set were closer in affinity, performance on all *OOD* tasks was high (Supplementary Fig.10). The challenging aspect of the “vs Weak” datasets may be due to the higher incidence of sequences where a single amino-acid substitution changes the binding class (Supplementary Fig.2a). Thus for *ID* and *OOD* performance, the distance between positive and negative datasets is important.

More generally, the difficulty of the *OOD* tasks varied across datasets due to differences in the similarity between positive and negative data, both in affinity and sequence space. A model trained on a simple task learns basic patterns but struggles to generalize to more complex tasks due to suboptimal decision boundaries. In contrast, a model trained on a complex task learns a more intricate decision space, allowing it to handle simpler tasks more effectively, as their patterns are often a subset of what it has already learned ^53^. In the future, it would be interesting to expand the definition of “similarity” or “task difficulty” to a multi-dimensional format not only dependent on affinity but also on developability ^17^ and sequence/structure motifs ^34^.

Lastly, we found that both *ID* and *OOD* performance were antigen-dependent. This is an observation we also made in another context (generative design of antibodies ^54^). This may be due to a variety of factors such as antigen shape, antigen-specific composition of training datasets, and epistatic considerations. As previously mentioned ^54^, this is beyond the scope of this work and requires further investigation in a separate study.

### Definition of binding rules

In this work, we define two types of “binding rules”: (1) total CDRH3 binding energy and (2) binding energies per amino acid residue in CDRH3 for a given antibody-antigen pair. The use of synthetic data provides a distinct advantage, particularly in studying per-residue binding energy rules. We can directly measure per-residue binding energies, rather than estimating them indirectly from total binding energies of mutational data. Knowing the binding energy of each residue allows for a more detailed downstream analysis compared to simply annotating residues as binders or non-binders ^34^. This approach contrasts with rules based on broader physicochemical properties such as charge, hydrophobicity, aromaticity of amino acids, and their distance to the target ^18,55,56^. We acknowledge that residues that do not contribute directly to binding energy may still influence binding indirectly through structural constraints. Therefore, we acknowledge that the rules we define here represent only a subset of the possible binding rules. In the future, it would be of interest to widen the definition of binding rules to interpositional interactions, also known as epistatic relationships ^57–59^, since we suspect that interpositional interactions are essential to identifying binding motifs.

### Negative data influence rule learning

We found that the composition of the negative dataset impacted the model’s ability to identify binding energies (rules) at both the sequence and residue levels (Fig.3). Specifically, (1) the correlation between output logits and antibody binding energy was particularly pronounced in the “vs Weak” and “vs Non-binder” tasks, (2) as was the correlation between individual amino acid contribution to binding energy and its contribution to model’s prediction (attribution). In contrast, for other tasks (“vs 1” and “vs 9”), the rule learning varied more substantially (Fig.3b,d). For several antigens, the learned rules were comparable, and the resulting attributions correlated well across all tasks (Supplementary Fig.8a). However, for most antigens, the amino acid importance information was confounded by the negative data, as evidenced by distorted attributions (i.e., a positive correlation with binding energy) in the “vs 1” and “vs 9” tasks (Fig.3d). This suggests that, while positive data alone may be sufficient to learn binding rules, negative data can either enhance or hinder rule learning.

A fundamental challenge is to fully decipher the information captured by model attributions. In our investigation, we did not observe scenarios where the energy contributions at the residue level and attributions exhibited correlations better than −0.6 to −0.7. Possible explanations include limitations in the attribution methods employed, incomplete energy computations (e.g., missed interactions by analyzing only CDRH3), and, more generally, the intrinsic complexity of per-residue contributions to binding. Hypothetically, the “vs Weak” task may enable the model to distinguish amino acids that are specific to the target interaction from those that are generally sticky, a distinction that may not be as apparent in other tasks. Interestingly, attributions obtained with DeepLIFT exhibited varying degrees of correlation with logistic regression weights (Supplementary Fig.7c). Attributions from the “vs Non-binder” task correlated more strongly with logistic regression weights, while those from the “vs Weak” task showed mixed correlations across antigens. This suggests that DeepLIFT captures some non-additive rules. However, due to its design, DeepLIFT does not provide direct information about feature interactions and therefore cannot be explicitly validated in this regard. This limitation highlights a broader challenge in the field—the need for methods capable of disentangling feature interactions. Beyond this, additional unanswered questions remain regarding attributions and their interpretation. For example, we note that the magnitude of attributions, evaluated by L2 Norm, was variable across task types, the attributions from “vs Non-binder” appearing to have consistently higher complexity (Supplementary Fig.8c,d).

A different perspective highlighting the complexity of rule learning in form of per-residue binding energies stems from the analysis based on epitope-based datasets, in which the positive class consists of only sequences binding to a single epitope as opposed to a single antigen (Supplementary Text 2, Supplementary Fig.9). Although the use of epitope-based datasets in comparison to the use of antigen-based datasets resulted in significant improvements in logits-based rule discovery, no significant and consistent difference was noted for the attribution-based rule discovery (Supplementary Fig.9d,e). Thus, although both per-sequence and per-residue energies appear to be discoverable, per-residue energies appear harder to disentangle.

### ML rule learning and prediction performance

In general, it is interesting to explore whether high-performing or more robust models rely on meaningful biological rules—capturing the properties that distinguish positive data from any negative. Specifically, we investigate whether rule learning covaries with performance as a function of target antigen and negative-data type used.

When analyzing how rule learning correlates with performance within tasks, i.e, across different target antigens, we observed that the total binding energy rule (logit rule) correlated with both *ID* and *OOD* performance (Fig.3c, Supplementary Fig.4b). This suggests that learning total binding energy partially explains variations in *ID* and *OOD* performance or that target antigen-specific features confound both performance and the learning of logit rules. In contrast, the per-residue energy rule (attribution-based rule) did not show correlation with *ID* performance within tasks (Fig.3e). Additionally, in tasks where this rule was learned most effectively (“vs Weak” and “vs Non-binder”), we did not see a strong connection between per-residue rule learning and *OOD* performance. Conversely, for models with substantial variation in per-residue rule learning (“vs 1” and “vs 9”), we observed a correlation with *OOD* performance. This suggests that residue-wise rule learning alone is not sufficient for high *ID* or *OOD* performance; however, if the learning of amino acid energy contributions is distorted by negative data selection, *OOD* performance can decrease substantially (Fig.3d, Supplementary Fig.4b).

Regarding observed biases towards specific antigens, we noted that both the SN10 and LR models tend to learn rules associated with antigens that are easier to predict, irrespective of how high-binders of an antigen are labeled as positive or negative. Specifically, we observed that reversing the roles of positive and negative datasets in ‘vs 1’ task can lead to a loss or identification of associations between energy and logits or attributions for positive data (Supplementary Fig.6, Supplementary Fig.7). Considering switching the role of positive and negative class in case of “vs 1” doesn’t impact prediction accuracy, it is noteworthy to highlight that in the case of “vs 1” the rule discovery for the positive class depends on which antigen serves as the positive class.

Examining the general relationship between rule learning and performance across different types of tasks, we found no clear dependence between rule learning and high prediction performance. For instance, models trained on “vs 9” and “vs 1” generally achieve higher *ID* performance than those trained on “vs Weak,” despite the latter learning binding rules more effectively across all antigens. This presents a counterexample to a hypothesis from a TCR-field study suggesting that models with higher performance are more likely to rely on biological rules ^60^. Our findings suggest that while *ID* performance may be necessary for rule discovery, it is not sufficient on its own. Instead, a model’s ability to learn detailed binding information represents an additional dimension of its overall performance. A slight refinement to this interpretation is that models with higher *OOD* performance are more likely to rely on robust biological rules. Despite our results showing that models trained on “vs Non-binder” and “vs Weak” exhibit similar levels of binding rule learning, *OOD* performance is lowest for “vs Non-binder” and highest for “vs Weak” among all tasks (Fig.2d, Supplementary Fig.3d). This discrepancy may arise from differences in how each task captures feature interactions. The “vs Weak” task appears to permit greater consideration of interactions between amino acid contributions, whereas the “vs Non-binder” task correlates more strongly with logistic regression weights across antigens—indicating a reliance on simple additive rules that align with an additive logistic regression model lacking interaction terms (Supplementary Fig.7).

Our results highlight the need for improved feature attribution methods and expanded binding rule definitions to more effectively reflect *OOD* robustness. Current attribution-based methods evaluate features independently, whereas feature interactions may play a critical role in biology. We suggest that *OOD* performance should serve as an orthogonal metric for benchmarking rule recovery methods, where a successful rule set and model interpretability approach will capture *OOD* robustness.

### Synthetic antibody-antigen binding data enables a detailed study of the impact of training dataset composition on ML performance

In this work, we used synthetic antibody-antigen binding data that was previously shown to help guide ML considerations on experimental data ^27^. The “Absolut!” simulation framework ^27^ was previously shown to have real-world relevance, accounting for many levels of complexity. It uses realistic 3D antigen sizes, shapes, and surface amino acid composition. The CDRH3 sequences used are derived from the murine repertoire and exhibit a physiological amino acid composition and positional dependencies. The antibody-antigen binding energy calculation is based on statistical potentials ^61^ derived from an analysis of known protein structures in the Protein Data Bank. “Absolut!” explores multiple structural conformations of CDRH3, thereby incorporating nonlinearity into sequence-to-affinity correspondence.

The use of synthetic data helped us explore how training data compositions impact ML performance, including the configuration of datasets by affinity. Although the affinity information allowed us to construct the datasets, we intentionally did not train the ML model to predict it. Our aim was to create an information bottleneck by designing the task as a binary classification, aligning with the most accessible experimental data for training immune receptor ML models. Current experimental data, as was used in this work, is only starting to become available in single sequence affinity-annotated format ^46,62^. However, these datasets may be generated with different technologies potentially compromising the comparability of affinity values. Furthermore, most publicly available datasets are not annotated or have affinity information with different cutoffs for binding/non-binding, which increases the influence of experimental batch effects and poses challenges for achieving robust prediction performance and meaningful rule discovery. Even if ignoring the mentioned limitations of current experimental data, the available experimental dataset size is one to two orders of magnitude lower (our synthetic dataset has ∼700k sequences in comparison to ∼25k for experimental) and the synthetic dataset covers more complexity, e.g., as evaluated by the higher number of covered antigens. The use of synthetic data enabled the testing of various sampling strategies and performance covariates (e.g., sequence composition, sequence entropy, antigen), which would not have been possible in that breadth with experimental data. Indeed, the testing of several hypotheses may be used for experimental design. In a sense, the “Absolut!” synthetic data can currently be seen as at least providing a lower boundary for the level of complexity of antibody-antigen binding datasets. Furthermore, synthetic data has the advantage of avoiding confirmation bias in explainability studies ^41,63,64^ as ground truth is available. Indeed, experimental data is distributed over an array of features, some relevant and others irrelevant for a given prediction task. These features may be structural properties, affinity, k-mer distributions, amino acid composition, and length of the CDRH3. It is generally not known in advance which features are irrelevant and, thus, which should be balanced in the training data (both positive and negative). Once identified, the irrelevant features could be balanced between the positive and negative datasets to prevent the model from learning spurious correlations. Towards this goal, here, as a first step, we explored this avenue by designing different negative classes. The large difference in performance among the different training data compositions supports the conclusion that there exist irrelevant features that are not equally distributed between different types of negatives, which can lead to high test set accuracy based on irrelevant features, a condition that leads to non-robust models with low generalization capabilities. In follow-up work, it may be interesting to investigate more granularly how the model behavior is impacted after certain features are balanced (e.g., forcing the distribution of accessible features such as k-mer distribution, resampling hidden features such as structural binding properties).

### Machine learning considerations

In this study, we focused on classification tasks because we mimicked the most commonly available datasets in immune-receptor field, which are often binary or semi-quantitative ^22,65,66^ (i.e., divided into categories based on binding strength). Furthermore, the negatives in the “vs 1” and “vs 9” datasets often are not annotated by affinity to a positive class target ^29,30,67^. Thus, we binarized our continuous affinity values to create semi-quantitative categories and evaluated whether these categories were sufficient for learning affinity properties, such as binding rules.

However, when affinity-annotated datasets are available, it may be feasible to adopt a two-step approach: first training a model as a classifier and then fine-tuning it on a smaller dataset with continuous affinity labels ^48,49^.

Building on this classification framework, we explored various ML predictive models. Specifically, we focused on the SN10 model because its architecture offers explainability and was previously identified as optimal for learning “Absolut!” data ^27^. Additionally, we tested a transformer-based architecture and incorporated language model embeddings; however, they did not yield better results for our tasks Supplementary Table 4, Supplementary Table 5). Our aim was to employ a neural network capable of accounting for interpositional dependencies. Consequently, we compared the SN10 model to logistic regression and demonstrated that SN10 outperforms logistic regression (Supplementary Fig.2c), suggesting that it can leverage non-additive sequence dependencies, particularly in the “vs Weak” task. Additionally, we observed that SN10 attributions showed varying correlations with logistic regression weights (Supplementary Fig.7), indicating differences in rule learning between these models. In the future, it would be of interest to study how rule learning depends on both input representation (sequence vs structure) and model architecture. Recent studies ^68,69^ investigating biases introduced by model architecture and input representation in predicting antibody mutational effects show that simpler models tend to learn binding interactions, whereas complex models capture broader biophysical properties related to protein folding.

We demonstrated that effective rule extraction relies not only on the mining of positive data but also on the careful selection of negative data. Our findings reveal that “vs Weak” and “vs Non-binder” establish decision boundaries based on features directly related to the binding phenomenon, such as total binding energy and per-amino acid energy contributions, notably both datasets containing sequences along a spectrum of binding energies across both the positive and negative parts. In contrast, “vs 9” and “vs 1” appear to focus more on spurious features. However, this hypothesis warrants further investigation.

Finally, in any kind of ML setting, information leakage (also called: data leakage) may occur where information from the testing or evaluation dataset inadvertently leaks into the training dataset. This may lead to overfitting, signifying that the ML model performs well on the evaluation data but fails to generalize to new unseen data. In biological sequence-based ML, it is therefore recommended to ensure that similar sequences do not appear in both the training and testing datasets ^1,70,71^. However, this approach does not account for the fact that very similar sequences may have entirely different functions (e.g., binding profiles), which is commonly observed in immune receptors ^65,72^ and which constitutes a great challenge. In our study, we did not observe a significant contribution to *ID* performance from sequences with the same label and 90% sequence identity between the training and testing sets. Sequences that are similar but have different targets or labels are challenging to predict and should therefore be distributed across train and test datasets (Supplementary Text 1, Supplementary Fig.2).

### Conclusion

By leveraging synthetic data, we identified dataset compositions with properties favorable for *OOD* generalizability and rule discovery, which we verified on independent experimental datasets. We observed that the closer the positive and negative training datasets are, the better the *OOD* performance of the model.

When constructing negative data, two main practical approaches exist: (1) subsampling existing nonspecific (negative) data and (2) generating negative data experimentally. Based on our findings, we recommend that when using the subsampling strategy, one can apply methods that select “hard” or “semi-hard” negatives from unlabeled data as discussed in Yang et al. ^73^. However, our work primarily advances the second approach—constructing negative data when there is control over the data generation process. In such cases, we recommend collecting data points that are close in terms of the metric of interest (e.g., binding affinity) to create a more challenging training set. It is important to note that while continuous measurements (e.g., binding affinity) are ideal, they are not always available on a large scale. In such cases, categorical labeling can be sufficient. For example, in flow-sorting experiments, antibodies are distributed into bins (e.g., high binders, weak binders, and non-binders). If such data are available or can be generated, we suggest using two adjacent bins (e.g., binders and weak binders) as the source of training data for machine learning models. In such a scenario, positive and negative data originate from the same experiment and the same sequence distribution. However, if positive and negative data are generated through separate experiments, experiment-specific biases might overshadow subtle differences between sequences of close binding affinity. Also, that might be another reason why “vs 1” and “vs 9”, negative datasets might not be reliable. Therefore, we advise caution when positive and negative data are generated through different experiments.

Additionally, deep mutational scanning (DMS) methods allow the creation of positive and negative datasets that are close in sequence and affinity ^65,74,75^ while providing information on epistatic interactions ^50^. Our work shows that DMS datasets will be crucial for teaching ML models sequence-to-function maps, where very small sequence differences can have large functional effects (Supplementary Fig.2).

Regarding rule discovery, we observed differences in learned binding rules depending on the negative dataset used for training. However, with current explainability methods, it remains difficult to disentangle individual amino acid contributions from their interactions (such as epistasis), which is critical for understanding protein structure ^50^ and function ^57,76^. To improve generalization across antigens, it will be important to identify binding motifs and develop a structured vocabulary for antibody-antigen interactions ^34,77^.

By demonstrating the added value of datasets with closely related positive and negative examples and the feasibility of learning binding energies solely from sequence data, we propose several avenues for further research. These include more detailed analyses of hard negatives and near-misses. Future work should also address the computational challenge of selecting negative data that are close to positive data ^14^, and developing more refined feature attribution methods and expanding binding rule definitions to better capture *OOD* robustness.

## Materials and Methods

### Synthetic antibody-antigen binding dataset generated using the antibody-antigen simulation framework Absolut!

We used a semi-synthetic dataset of antibody-antigen binding as previously generated by Robert et al. and Akbar et al. ^27,54^. Briefly, this dataset comprises real-world antigen and antibody (CDRH3) proteins with antibody sequences from mouse repertoires ^78^ and with the antigens’ 3D structures sourced from the Protein Data Bank (PDB) and transformed into a 3D lattice format. The authors of the dataset applied an exhaustive docking method to evaluate the binding of discretized rigid antigens with flexible CDRH3s of length 11 from the antibody side. The result included the acquisition of binding energy values, binding positions that minimize free energy, and the identification of interacting amino acids between antigens (epitope) and CDRH3s (paratope). Each antigen was associated with a set of 11-mers derived from CDRH3s and ranked according to their binding energy. Those in the lowest 1% percentile, with an average of 51,814.7 sequences, were labeled as high-affinity binders (or simply ‘binders’). The 1-5% percentile, averaging 205,069.3 sequences, were labeled as weak-affinity binders. Finally, the highest 95% percentile, with an average of 391,498.8 sequences, were labeled as non-binders. This labeling scheme aligns with the methodology used in the study by Robert et al. ^27^. For our study, we used antibody binding data from 10 antigens, with PDB-IDs 3VRL, 1NSN, 3RAJ, 5E94, 1H0D, 1WEJ, 1ADQ, 1FBI, 2YPV, 1OB1.

### Prediction tasks, training, and test datasets

We used balanced CDRH3 datasets (30k antibody sequences for training [15k positive class, 15k negative class], 10k antibody sequences for testing [5k positive class, 5k negative class]), which varied by the positive class antigen and the negative dataset type (Table 1). The positive class of the dataset consisted of binder sequences, i.e., sequences having binding energies between 0–1% percentiles.

The negative class of each dataset was built depending on the binding prediction task type, as follows:

● vs 1 – sequences that are binders for another antigen but not the positive class antigen.
● vs 9 – sequences that bind to one of the 9 other antigens and do not bind to the positive class antigen, represented in equal proportions. To illustrate, in the training set of a total of 15,000 sequences, each of the 9 antigens has approximately 1,666 (+/- 1) associated sequences (∼15,000/9).
● vs Weak – sequences that bind the positive class antigen, but with binding energies in the 1–5% percentile.
● vs Non-binder – sequences with binding energies in the >5% percentile for the positive class antigen.

We generated replicates to ensure the reproducibility of the experiments. Specifically, six different train-test splits were generated for each prediction task and positive antigen. For one of the splits, we trained and evaluated the model four times with various seeds, therefore controlling for the randomness in model initialization and training. The positive class of the datasets is kept constant across the tasks and in each replicate experiment. For example, the positive class data in *1ADQ Binder* “vs Weak” is the same as the positive class data in *1ADQ Binder* “vs 9”. The sequences corresponding to each positive and negative dataset were randomly sampled to achieve the previously mentioned dataset sizes. Throughout our work, we employed negative controls by randomly shuffling the class labels of sequences for each task. The prediction performance was evaluated on hold-out test datasets using accuracy metrics, calculated as the number of correct predictions divided by the total number of predictions. The *out-of-distribution (OOD)* performance was evaluated by calculating the performance on the test sets of other tasks, e.g. “vs 9” performance on a “vs Weak” test set. The out-of-distribution estimations were performed over the same set of replicates as in the in-distribution case.

### Machine learning model architecture, training, and evaluation

We trained a simple neural network, previously ^27^ and henceforth called SN10 ^27^, for binding prediction tasks using CDRH3 sequences. The sequences, each consisting of 11 amino acids, are one-hot encoded, resulting in a dimensionality of 220. The neural network architecture consisted of 220 neurons in the input layer, 10 neurons in the hidden layer (ReLU as activation), and 1 output neuron (sigmoidal activation). For training, after exploration of the parameter space, we used binary cross-entropy loss with the Adam optimizer with a batch size of 64, a learning rate of 0.001, and a momentum of 0.9, with no weight decay. We trained the models for 50 epochs and inspected the loss curves, which did not show signs of overfitting (Supplementary Fig.12). To further enhance the model’s performance, we implemented the technique of stochastic weight averaging (SWA) ^79,80^, which involves averaging multiple sets of weights obtained during the training process to reduce the variance and improve the generalization performance of the model. We apply SWA after epoch 3/50, using the SWA implementation available in Pytorch v1.12.1 ^81^. A logistic regression-based model was further implemented for binary classification as a baseline for comparison. The model was trained and evaluated on the same datasets using one-hot encoded CDRH3 sequences, as previously described for the SN10 ^27^.

We tested additional more complex architectures in the process of evaluating the use SN10 as the main model of our study. The Transformer model architecture consisted of an encoder with a dimensionality of 24, a vocabulary size of 30, 4 attention heads, and a single feed-forward layer with a dimensionality of 128. It incorporated a 0.1 dropout rate and used ReLU as the activation function. The model was trained on a subset of the datasets used for SN10 evaluation, utilizing the Adam optimizer (momentum 0.9, weight decay 0.0) with a batch size of 128 and a learning rate of 1e-6 for 500 epochs. The architecture was implemented using PyTorch’s transformer building blocks, with the code available on GitHub. The employed training loop was identical to that used for SN10. For the PLM-based SN10, we utilized the ESM2b protein-language model ^82^ to generate 1280-dimensional embeddings of the CDR3Hs, as well as AntiBERTa2 (referred to as Antiberta2) to generate 1024-dimensional embeddings ^83,84^. Implementation wise the EmbedAIRR tool developed by our lab was utilized (https://github.com/csi-greifflab/embedairr). These embeddings were used as inputs to the SN10, transforming its architecture into a feed-forward neural network with an input dimensionality of 1280, 10 hidden units, and 1 output unit, last 2 layers as in the original SN10 design. After confirming that the training protocol for SN10 did not cause overfitting for the PLM-based SN10 models, we applied the same training protocol as described above. Summarized results based on transformers and PLM-based SN10 are presented in Supplementary Table 4 and Supplementary Table 5.

### Information theory-based calculations

Jensen-Shannon distance (JSD) is a measure of dissimilarity between two probability distributions. It is derived from the Kullback–Leibler divergence (KL) and calculated using the following formula:

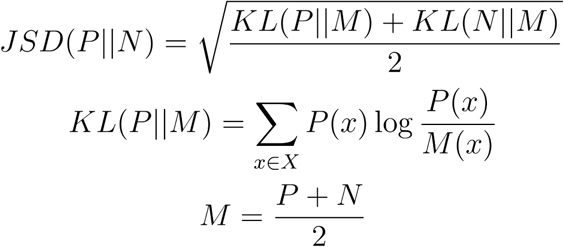

In our study, P and N represent the probability distributions for the positive and negative data sequences, respectively. The JSD was computed using the scipy.special.distance package. As a measure of the total differences in marginal distributions for positive versus negative data sequences (i.e., excluding differences in statistical interactions), we utilized position weight matrices (PWMs) that represent the probabilities of each amino acid occurring at specific positions within binder or non-binder sequences. The JSD in our setting is the sum of JSDs across each position. PWMs were calculated using the bio.motifs package from Biopython ^85^.

Sequence Shannon entropy calculations were performed to measure antibody sequence diversity for given sets of sequences (e.g. 5E94 binders, 3VRL weak binders) (see Supplementary Fig.1). The Shannon entropy was computed using the scipy.stats Python module to quantify the information content based on the PWM (computed as previously described) of the set of sequences.

### Attribution analysis

To better understand the rules learned by a SN10, we employed interpretability techniques and applied them to test set examples to gauge the importance of input features for each sequence, importance in this case meaning contribution to the prediction. We used the DeepLIFT ^45^ implementation from the Captum library v0.5.0 ^86^ as a fast method that approximates Integrated Gradients ^87^. The computation requires baseline sequences, which were obtained by randomly reordering input sequences, a technique known in the literature as the shuffling approach ^88^. Although using a zero input vector as a baseline is common and plausible in certain applications, in the case of sequences, it is problematic since a zero input vector does not represent any sequence, i.e., there is no sequence that would be one-hot encoded as a zero input vector. The shuffling approach is based on random sequences and has been the preferred approach for biological sequences in the recent literature ^88,89^. We generated ten shuffled versions for each CDRH3 sequence and ran DeepLIFT ten times per sequence, once for each of the shuffled baselines. The final attribution values were obtained by averaging the results from these ten runs. The attributions were estimated with respect to the predicted logits, not expits. We used local attributions ^90^, since we were interested in the impact to the prediction of each amino acid in each sequence. The implementation of this calculation in Captum ^86^ involved setting *multiply_by_inputs* to True. Attribution analysis was applied to synthetic and experimental data. However, since there was no binding energy data per amino acid in the experimental dataset, we could compare attributions to ground truth per-residue energies only in the synthetic dataset. Attributions were computed across all replicates except for “vs 1”, where due to the much larger number of datasets, we restricted the computation to once per dataset.

### Epitope-based analysis

For the epitope-based analysis, we generated new datasets while restricting, in each case, the positive datasets to consist only of sequences that bind to the major, i.e. most frequent, epitope of an antigen. For this analysis we selected the antigens that contain a sufficient number of epitope-specific sequences to allow for a training set size with the same size as in the antigen-based analysis (N_train_ = 30000). However, the test set size, due to the lower availability of sequences, had to be reduced w.r.t. antigen-based analysis (N_test_ = 6000). Due to the training size restriction we had to limit the epitope-based analysis to 3 antigens: 1H0D, 1WEJ, 1OB1. In all our workflows and results, to distinguish between antigen- and epitope-based, we depicted the epitope-based datasets with the same antigen id, followed by “E1” (indicating the most frequent epitope): 1H0DE1, 1WEJE1, 1OB1E1. To generate the training and test sets, we filtered and adjusted the positive datasets by keeping the sequences that bind only to the major epitope and kept the same negative datasets as for the corresponding antigen-based analyses. After generating the corresponding datasets, we performed identical computations as for the antigen-based analyses that were shown in the main text.

### Absolut! binding energy calculations

We refer the reader to Absolut! ^27^ for a detailed explanation of the computation of binding energies. Briefly, in the 3D lattice, only non-covalently bound neighboring amino acids are considered to interact. The binding energies are then computed by summing the Miyazawa-Jernigan potential for each interacting pair of residues. By summing the Miyazawa-Jernigan potential for all interactions in which a CDRH3 residue is involved, the energy contribution of that residue to CDRH3:antigen binding energy is computed.

### Experimental antibody-antigen binding datasets

#### The experimental dataset

We identified a publicly available experimental dataset ^46^, generated for high-throughput antibody discovery using next-generation sequencing, ribosome display and affinity screening techniques, referred to throughout the text as the “Porebski dataset” or “Experimental dataset”. The technique generates binding kinetics data for an antibody library, which can be used to derive measures of antibody affinity and thus to perform screening. Fluorescently-labeled antigen accumulates around higher-affinity binders and images acquired at different antigen concentrations are processed to derive a per-sequence spatially integrated signal based on the fluorescence intensity, which is associated with affinity. The authors of the study performed screening of 2 anti-HER2 antibody libraries, one library (HER2affmat) generated through affinity maturation of G98A, a low-affinity anti-HER2 scFv, and another library (HER2mllib) with higher-affinity antibodies that was generated using a protein language model trained on the HER2affmat. To operate on higher affinity antibodies, we used the HER2mllib.

#### Processing

We kept the unique sequences that had ≥ 12 replicates and that contained no unresolved amino acids and no stop codons, resulting in 24,790 sequences. All CDRH3 sequences had a length of 21 amino acids. Leveraging the results of the study, we used the *intensity 8* measure (the fluorescence-derived signal at a specific antigen concentration from the study) as a proxy for affinity (higher intensity ∼ higher affinity), as it has been used in the original paper to classify antibodies based on affinity. Using two thresholds, we classified the sequences in *Binders* (*intensity 8* > 150, 2638 sequences, 10%), *Weak* (122 < *intensity 8* < 150, 3574 sequences, 15%) and *Non-binders* (*intensity 8* < 122, 18578 sequences, 75%). We used thresholds with different percentiles compared to Absolut! to adapt to the smaller dataset size and obtain more sequences per antibody class for more reliable training and testing. Based on this classification, we built “vs Weak” and “vs Non-binder” datasets. In addition, we included a negative dataset consisting of randomly reordered CDRH3 sequences from the Binders – “vs Randomized negative”.

#### Model training and attributions

We trained SN10 models with replicates and computed attributions using exactly the same methodology as described above for the Absolut! dataset by adapting the input size to 21 amino acids and the size of the training set (N=4000) and of the test set (N=1000). Inspection of train and test loss by epochs showed no signs of train set overfitting in our setup. All settings, including performance metrics, were identical to those used with synthetic data.

## Code availability

The code is available at https://github.com/csi-greifflab/negative-class-optimization.

## Data availability

All synthetic data used in this work stems from the “Absolut!” database, available on https://greifflab.org/Absolut/^27^. We make all the processed datasets, trained models, attribution values, and results available at https://doi.org/10.5281/zenodo.11191740. The experimental Porebski dataset is available at https://doi.org/10.5281/zenodo.8241732.

## Acknowledgments

None.

